# Universal kinetics for engagement of mechanosensing pathways in cell adhesion

**DOI:** 10.1101/292367

**Authors:** Eugene Terentjev, Samuel Bell, Anna-Lena Redmann

## Abstract

When plated onto substrates, cell morphology and even stem cell differentiation are influenced by the stiffness of their environment. Stiffer substrates give strongly spread (eventually polarized) cells with strong focal adhesions, and stress fibers; very soft substrates give a less developed cytoskeleton, and much lower cell spreading. The kinetics of this process of cell spreading is studied extensively, and important universal relationships are established on how the cell area grows with time. Here we study the population dynamics of spreading cells, investigating the characteristic processes involved in cell response to the substrate. We show that unlike the individual cell morphology, this population dynamics does not depend on the substrate stiffness. Instead, a strong activation temperature dependence is observed. Different cell lines on different substrates all have long-time statistics controlled by the thermal activation over a single energy barrier ∆*G* ≈ 19 kcal/mol, while the early-time kinetics follows a power law ~ *t*^5^. This implies that the rate of spreading depends on an internal process of adhesion-mechanosensing complex assembly and activation: the operational complex must have 5 component proteins, and the last process in the sequence (which we believe is the activation of focal adhesion kinase) is controlled by the binding energy ∆*G*.

## 1 Introduction

Matrix stiffness is known to affect cell size and morphology (1, 2). When cells are plated onto soft substrates, their footprint will not increase as much as on stiff substrates, and their spreading will be more isotropic: resulting cells will be round and dome-like in shape. On stiff substrates, the same cells will spread very strongly, develop concentrated focal adhesion clusters and stress fibers of bundled F-actin, and eventually polarize to initiate migration. This leads to several well-documented biological functions in tissues: variable stem-cell differentiation pathways (1, 3), the fibroblast-myofibroblast transition near scar tissue (4–6), fibrosis in smooth-muscle cells near rigid plaque or scar tissue (7, 8), and the stiffer nature of tumor cells (9, 10). The definitive review (11) summarizes this topic. There are several different mechanosensing processes acting either simultaneously, or in different circumstances. One of the key physical mechanisms of stiffness sensing is thought to be based on the extracellular latent TGF-*β* complex (5, 12); another key mechanism involves the intracellular complex incorporating integrin and focal adhesion kinase (FAK) (13–15). There is also a lot of discussion of integrin itself, or vinculin, using their catch-bond characteristics to produce mechanosensing (16, 17), however, we would argue that these proteins do not possess a catalytic domain and therefore cannot produce a required chemical signal: the output of a mechanosensor (15). It is important to note that many catch-bond models use the collective activity of many integrins in mature focal adhesion clusters, while in this work we focus on the properties of individual adhesion-mechanosensing protein complexes. This is more important in the initiation of spreading, where mature clusters are yet to develop. We also emphasize that here, and in the rest of this paper, we are discussing isolated cells on a substrate: when cells adhere to each other, their shape transitions are controlled by other mechanisms, based on cadherin and associated pathways (18).

The dynamics of cells spreading has been studied extensively, and several characterstic universal features have been established (11, 19, 20). In particular, the average cell area has been shown to grow with time as a power law, often with the radius of cell footprint being *R ∝ t*^1/2^ (21–23). Several mechanistic models of how the cell spreading is achieved after the adhesion to extracellular matrix (ECM) is established (19, 21, 23). However, these papers deal with the characteristic rate of spreading for individual cells, and so neglect stochastic effects in the population dynamics. Here, we observe population dynamics directly, and account for variability in the spreading behavior over different cells.

While reporting and discussing the cell area increase on stiffer substrates, the seminal paper (2) also presents some data on the time-dependence of cell spreading, which already gives a hint for our central experimental finding: the time for cells to engage their mechanosensing pathways, and spread, does not depend on the substrate. In this paper we investigate the time-dependence (kinetics) of mechanosensing response, asking the question: how long does it take for the cell to recognize the nature of its substrate, and respond by engaging the signaling pathways and enacting the required morphological change (spreading on the substrate)? Figure 1 illustrates the point: plots (a) and (b) show the same cells: immediately after planting on the substrate, and after some time, when several cells have already responded by engaging their spreading. We plated two very different cell lines (NIH/3T3 fibroblasts and EA.hy927 endothelial cells) on a variety of substrates that span the range of stiffness from 30 GPa (stiff glass) to 460 Pa (very soft gel), registering the characteristic time at which the initially deposited planktonic cells engage their mechanosensing response.

**Figure 1:**
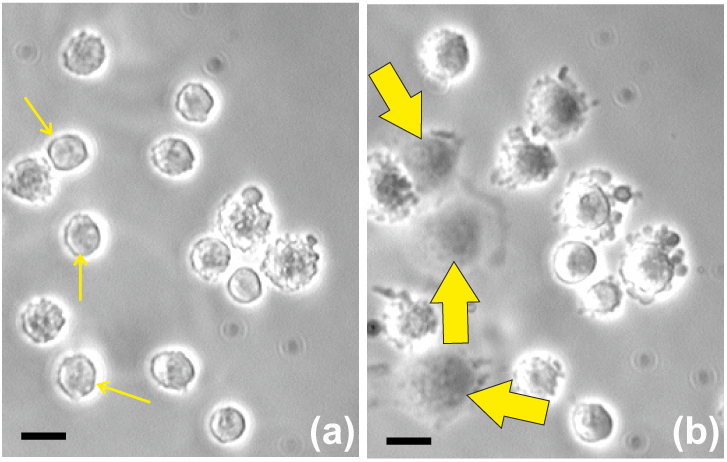
Illustration of mechanosensing response and cell spreading. Plots (a) and (b) show the same cells: immediately after planting on the substrate (solid glass with fibronectin), and 15 min later, when several cells have already responded by spreading (labeled by mathcing arrows). Scale bar = 20 *µ*m.

We discover three remarkable and unexpected facts: [1] the rate of mechanosensing response is completely universal, not depending on the stiffness of substrates (in contrast to the final cell morphology, which strongly depends on it); [2] the rate-limiting process in mechanosensing response is the same in all cells, and is the activation of FAK (with the energy barrier ∆*G* ≈ 19 kcal/mol in good agreement with simulation studies); [3] the onset of mechanosensing response is controlled by the kinetics of the adhesion-sensor complex assembly, its universal power-law dependence *t*^5^ suggests that there are exactly four steps in the complex assembly, i.e. five constituting proteins (followed by the FAK activation step of sensor firing). We also measure the sum of the binding energies of proteins in this assembly process, and find that this is non-universal: it depends on the cell type and probably affected by their biological function.

## 2 Materials and Methods

### Cells and cell culture procedures

We chose to study endothelial cells and fibroblasts because their adhesion behaviour is important for understanding cardiovascular diseases and tissue engineering. There are different types of endothelial and fibroblast cells available. Primary cells are directly taken from donor tissue and then grown in cell culture conditions. They can be grown in culture for a specific amount of time before they undergo senescence and die. The advantage of primary cells is that they are as close to in-vivo cells as possible, but as they are taken from different donors, their behaviour is less reproducible. Immortalized cell lines are obtained from primary cells by, for example, transfection or fusion. This results in a change in their DNA, leading to indefinite proliferation. This makes it easier to handle them in multiple long-term experiments, and makes such experiments more reproducible, but at the same time many immortalized cells have some tumorous behavior (24, 25). We choose to use immortalized cell lines: NIH/3T3 murine fibroblasts (obtained from ATCC) and EA.hy927 endothelial cells.

NIH/3T3 fibroblasts are very well characterized, as they have been used in many cell studies since their establishment as cell line; they have also been used in cell adhesion studies, making them a good choice for our experiments (26, 27). EA.hy927 is a cell line established in 1983 by the fusion of HUVEC with a lung carcinoma line (28). It has since become a widely used and thus well characterized cell line, popular in studies of cardiovascular diseases. EA.hy927 have also been used for adhesion strength assays (29).

Cells were normally cultured at 37C and 5% CO2 in Dulbecco’s modified Eagle’s medium (Greiner) with 10% fetal bovine serum and 1% Pen/Strep (solution stabilized, with 10,000 units penicillin and 10 mg streptomycin/mL), from Sigma Aldrich (this standard medium is abbreviated as DMEM). For a comparative study of the role of nutrient in the medium, we also used phosphate-buffered saline (PBS), from Thermo Fisher Scientific during the spreading experiments. Cells were subcultured in DMEM every 3 days, at about 70% confluency, by trypsinization, to avoid the formation of big lumps of cells, thus ensuring that we maintain a single cell suspension. Cells were trypsinized for 5 min (Trypsin-EDTA 0.05%). The solution was then neutralized by added complete growth medium and centrifuged at 1000 rpm for 5 min.

The use of Pen-Strep can be questioned. Antibiotics have been used prophylactically to prevent bacterial infections in cell culture for many years, and they are still being used. It was the introduction of antibiotics that allowed the widespread development of cell culture methods in the first place, as bacterial contamination was a major problem (30). However, although toxicity experiments found that antibiotics were harmless to mammalian cells (31), there are concerns about the use of antibiotics in cell culture associated Biophysical Journal 00(00) 1–10 with a neglect of aseptic technique and possible side effects of antibiotics. Many adhesion strength studies use Pen/Strep or other antimycotic or antibiotic solutions in the cell culture, and we followed this procedure as well. We have tested our results on several parallel cell cultures that did not use Pen/Strep, and confirmed that no significant difference was inflicted on our results.

### Substrates of varying stiffness

To span a wide range of substrate stiffness, we used standard laboratory glass (elastic modulus 30 GPa), and several versions of siloxane elastomers: Sylgard 184 and Sylgard 527, the latter used with the compound/hardener ratio of 1:1 and 5:4. The resulting elastomers were tested on a standard laboratory rheometer (Anton Paar), giving the values of equilibrium modulus *G* = 460 Pa for (S527 5:4), 480 kPa (for S184), and 30 GPa for glass (zero-frequency limit shown in the Supplementary plot, Fig. S1). For comparison, the stiffness of typical mammalian tissues is commonly reported as: 100 Pa – 1 kPa in brain tissue; ~3 kPa in adipose tissue; 10 – 20 kPa in muscle; 30 – 50 kPa in fibrose tissue; up to a few MPa for bone. We avoided applying the commonly used plasma treatment, as this was making the surface highly uneven on a micron scale, which would affect the adhesion. All surfaces were cleaned by ultrasonication in 96% ethanol for 15 min, and then incubated with 10 µg/mL fibronectin in PBS for 45 min.

### Experimental procedure and data acquisition

In our standard cell-spreading experiment, the cell culture dish was inserted into a closed chamber that maintained controlled temperature with an active water bath, and the CO_2_ atmosphere, while allowing a microscope observation from the top. The cell culture (density 5 × 10^5^ cells per ml, counted by the Neubauer chamber) was placed over the entire substrate. Cells were left to adhere to the substrate for 2 min, at which point the culture dish containing the substrate is filled slowly with fresh medium to reduce the cell density. This was to prevent new cells depositing, and cell clusters forming on the substrate. Only the cells attached to the substrate at this point were included into the subsequent counting.

To obtain a population distribution of the onset time of cell spreading, we had to choose a spreading criterion, which would be clear and easily distinguishable to avoid counting errors. We choose to count the initial onset of visible spreading, seen as the transition between the near-spherical cell initially planted (physically attached) on the substrate, and the cell with adhesion processes engaged and its shape developing an inflection zone around the rim (see Supplementary Fig. S2 for illustration). This turns out to be easily identified as the near-spherical cell has the sharp edge, and also a lensing effect of focusing light, which disappears on the transition to a more flattened shape. A schematic of this can be seen in Fig. S3. It must be emphasized, that in order for our cell counting to be meaningful, the cells have to be isolated on the substrate: once the cells come into contact with each other, many other mechanosensing mechanisms engage (based on cadherins, and other cell-adhesion systems), and they spread much more readily and more significantly. That is why our initial cell density was chosen such that the initial attachment is in isolation, and our spreading criterion is applied before they spread sufficiently to come in contact (as some cells in Fig. 1 have done).

In each individual experiment (given substrate, fixed temperature, and other parameters), once the cells were deposited on the substrate, and the clock started, we took broad-field microscopic images at regular time intervals, and counted the fraction of cells that have crossed the threshold defined by our spreading criterion. This produced a characteristic sigmoidal curve for each experiment (see Fig. 2): the fraction of cells ‘engaged in spreading’ starting from zero at *t* = 0 and saturating at near-100% at very long time (if we exclude the occasional cell mortality). The typical sample size was 100-120 cells in each experiment (field of view). The main sources of error were: inconsistency of application of the spreading criterion in image analysis, imperfections of fibronectin coverage on substrate, temperature fluctuations, and of course the natural cell variability. All of these are random errors, with no systematic drift. We were satisfied that the results were reproducible, and errors did not dominate the data trends.

**Figure 2:**
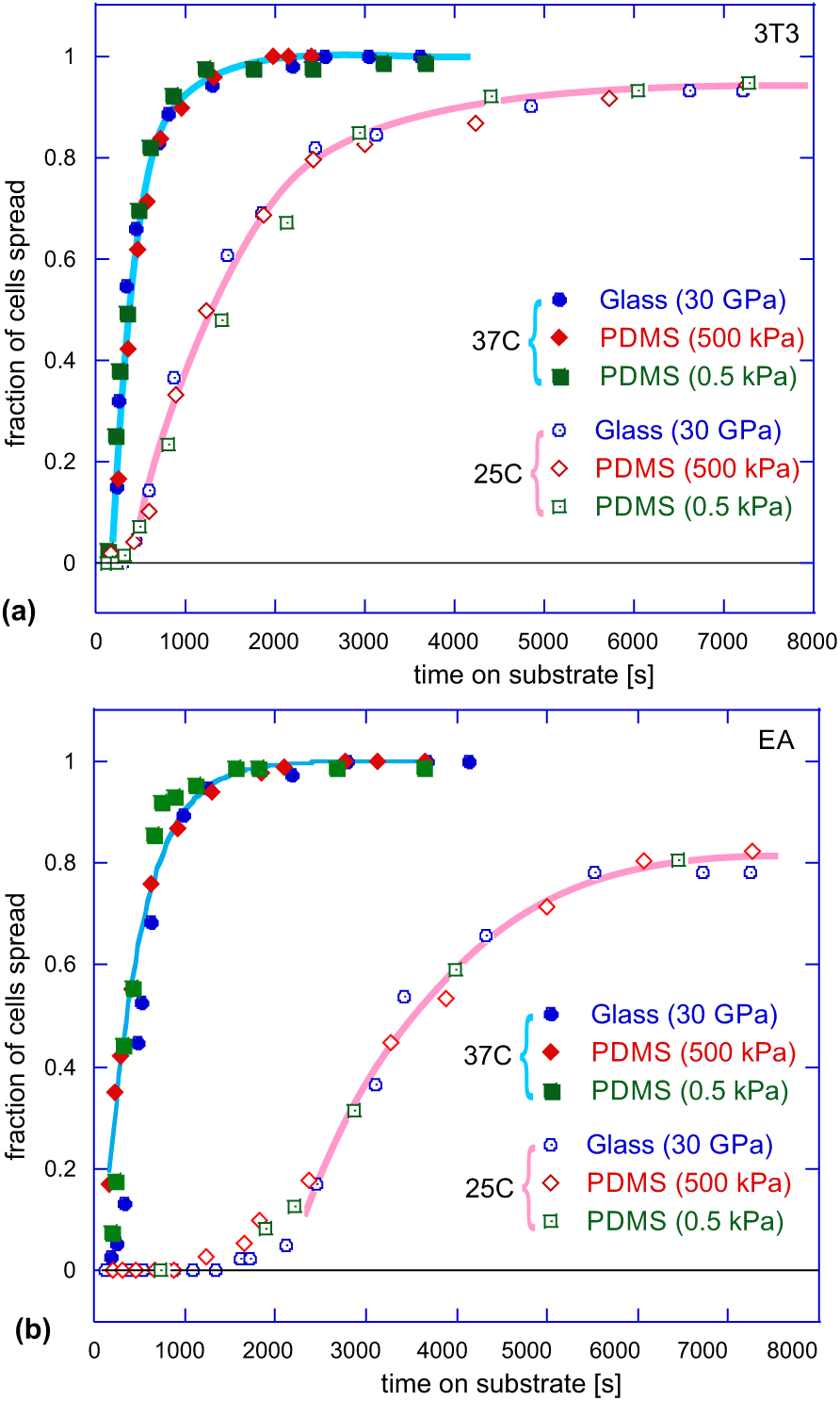
Cumulative population dynamics of cell spreading. Plots (a) and (b) show the growing fraction of cells engaged in spreading on substrates with different stiffness for 3T3 fibroblasts and EA endothelial cells at two different temperatures each. It is clear that the dynamics is not affected by the substrate stiffness, but changes with temperature. In the remainder of this paper, we analyze in detail the long-time behavior of these cumulative curves as they approach saturation, and the behavior at short times when the onset of mechanosensing response occurs.

## 3 Results

It is well-established that cells spread differently on substrates of differing stiffness (there is also a strong dependence on the ECM protein coverage (32), but this was not a variable in our study). The area of a spread cell is invariably greater on stiff substrates, and the shape has different features: more round and symmetric on soft substrates and highly asymmetric, with large strong focal adhesions on stiff substrates (the key article by Janmey et al. (2) makes a very detailed study of this effect). We also see the same effect, for both our cell types, on all substrates.

The first, unexpected, result is presented in Figure 2. These two plots illustrate the cumulative population dynamics: after planting, cells spend several minutes making a decision before starting their spreading – to a wide-area footprint on stiff substrates, or to a more round dome-shape on soft substrates, in agreement with classical studies (1–3) – while the timing of cell spreading is completely insensitive to the substrate nature. The kinetics of mechanosensing response is exactly the same on each substrate; the point of steepest gradient in the cumulative curves in Fig. 2 marks the most probable time for the onset of cell spreading. The work of Sheetz et al. (33) has reported a similar effect (the rate of spreading did not depend on the degree of ECM protein coverage on the surface). The second remarkable fact revealed in Fig. 2 is that the kinetics of cell spreading is strongly affected by temperature.

This leads us to the main conclusion of this paper: the rate of mechanosensing response is an internal cell characteristic, determined by the nature of its sensor and the signaling pathways that translate the sensor output into the morphological response of the cell. The magnitude of this response is affected by the sensor signal strength – but the timing of this process is determined by the internal cell organization, and its thermally-activated processes.

### Long-time trend: rate limit of mechanosensing

To examine the effect of temperature in greater detail, in Fig. 3 (a,b) we plotted the same cumulative spreading-dynamics curves for the two cell types on glass (as we are now assured that these curves are the same on all substrates). It is noticeable that the initial lag is greater in the EA cells, and that at low temperature the saturation level drops significantly below 100% – presumably because more cells disengage (or die) at low temperature, reducing the saturation fraction. The same effect is much enhanced for the the nutrient-starved cells in the PBS medium, see in Fig. 3(a): the kinetics of mechanosensing response is very slow in this case, and a large fraction of cells do not engage at all, which is not surprising because the ATP-consuming processes are involved in generating the cytoskeletal force and in converting Rho GTPases. But the generic shape of the cumulative curve is universal, and the random spread of data within each individual experiment is not excessive. We then look to analyze the trends in this time dependence.

**Figure 3:**
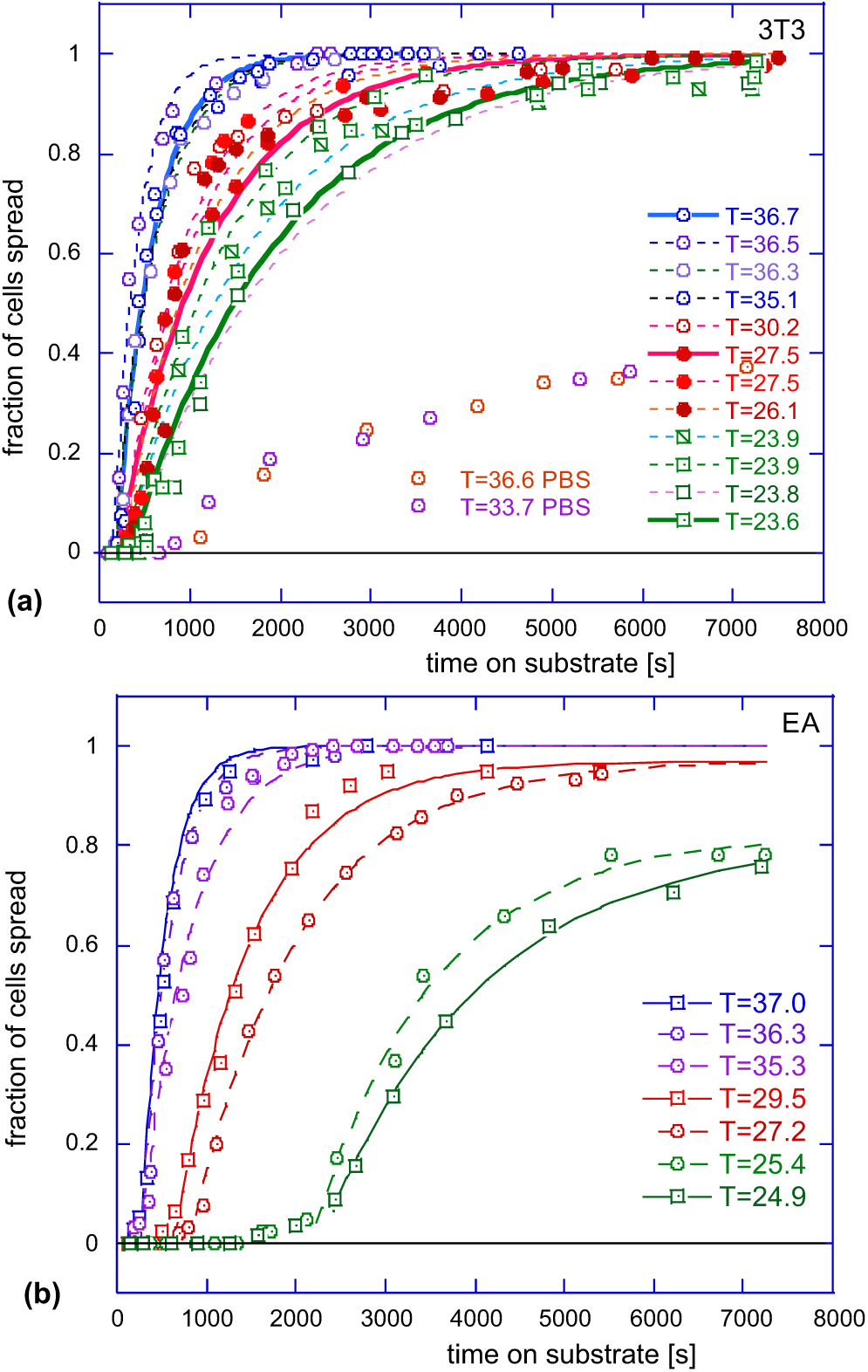
Cumulative population dynamics of cell spreading. Plots (a) and (b) show fraction of spreading cells on glass, at many different temperatures; for 3T3 fibroblasts and EA endothelial cells. Lines in all plots are the fits of the long-time portion of data with the exponential relaxation curves, producing the fitted values of the longest relaxation time *τ* (see text).

The curves of the generic shape seen in Figs. 2 and 3 are encountered in many areas of science, and their characteristic foot at early times, especially obvious at lower temperatures, is usually associated with a lag in the corresponding process. We will discuss this early-time regime separately, later in the paper, but first we fit exponential relaxation curves to the long-time portion of the data (as the fit lines in Fig. 3 indicate): *Q*(*t*) = *A* (1 exp[(*t t*_lag_)*/τ*]. The Supplementary Information gives the table of values of *A* and *τ* for each curve, but it is clear from the plots that the fitting to the single-exponential relaxation law, with just two parameters since *A* is known for each curve, is very successful. The characteristic relaxation time *τ* markedly increases at low temperatures. It is interesting that such a characteristic time associated with the ‘spreading of an average cell’ has been discussed in (19), giving the same order of magnitude (of the order of magnitude 50-100s).

To better understand this dependence on temperature, we tested a hypothesis that this relaxation time is determined by the thermally-activated law by producing the characteristic Arrhenius plots of relaxation times, for both cell types, see Figure 4. It is remarkable that both cells show almost exactly the same trend of their relaxation time: the rate limiting process in their mechanosensing pathways is the same: *τ* = τ_0_*e*^*∆G/k*_*B*_ *T*^, with the activation energy *∆G ≈* 18.7 *±* 1.5 kcal/mol, and the thermal rate of attempts 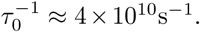 Both values are very sensible: this magnitude of ∆*G* is typical for the non-covalent bonding energy between protein domains (34), and this rate of thermal collisions is in excellent agreement with the basic Brownian motion values.

**Figure 4:**
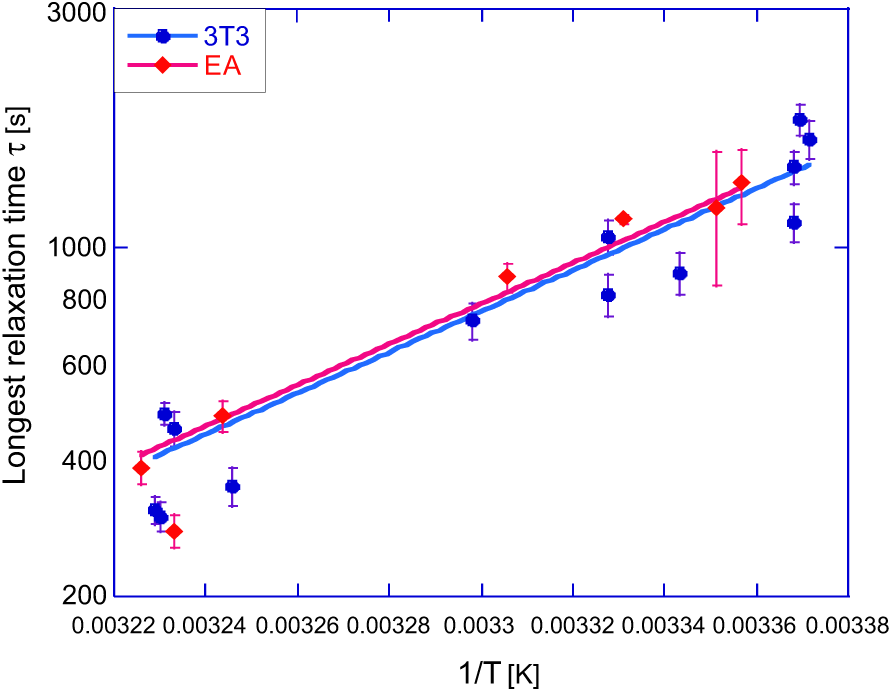
The Arrhenius plot of the longest relaxation time (log(*τ*) vs. inverse absolute temperature) from the exponential fits in Fig. 3 (a,b), giving the same value of binding energy ∆*G ≈* 19 kcal/mol, for both types of cells.

### Early-time trend: rate of complex assembly

After discovering that the late-times (rate-limiting) dynamics of mechanosensing response is quite universal, across different cells and substrates, it becomes clear that the marked difference between different curves in Fig. 3 lies in the early-time behavior: something that we have called a ‘lag’ following many similar situations in protein self-assembly. To examine this early-time regime more carefully, we re-plotted the same time series data on the log-log scale in Fig. 5 (a,b). First of all, from examining these plots it is clear that there is no such thing as a lag time: the log-log plotting reveals that the process is active from the very beginning (*t* = 0) and the plotted value grows as a power-law of time. The only reason that we appear to see a ‘lag’ is because our experimental technique of counting the cells engaging in spreading did not permit values below 0.01 (1%) to be resolved in this plot; the same certainly applies to other experimental situations reporting similar kinetic data. The trend illustrated in Fig. 5 is clear: the early onset of cell spreading follows the universal power law, and the fitting of all our data sets gives *Q*(*t*) = α*t*^5^ with very good accuracy, where only the prefactor *α* depends on temperature and the cell type. We find this result truly remarkable: similarly to the universal value of binding energy that controls thermally-activated rate-limiting relaxation time *τ*, this very specific *t*^5^ power law appears to be the only sensible fit of the early-time data for different cells, temperatures, and substrates.

**Figure 5:**
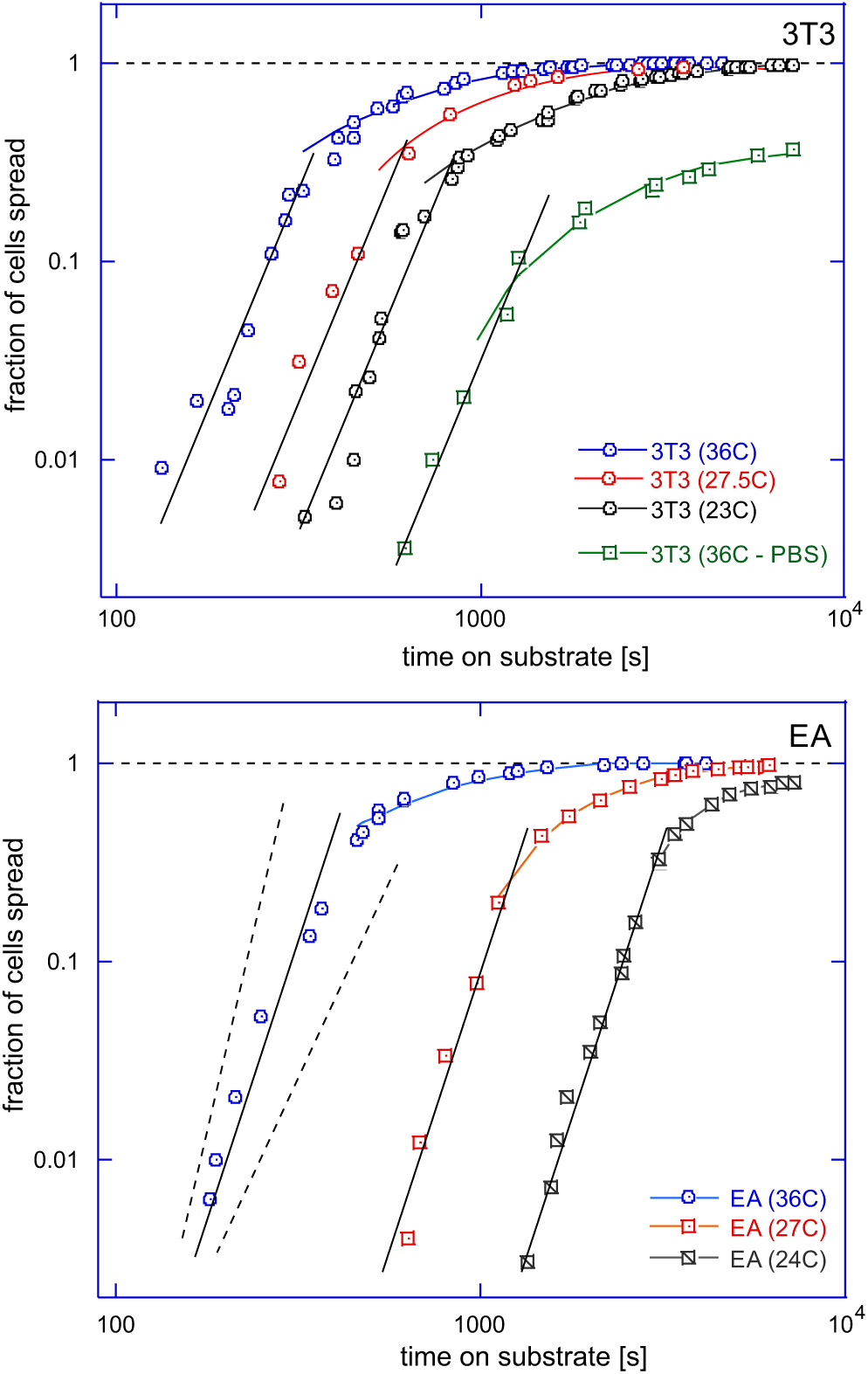
Analysis of the short-time dynamics of cell spreading. Plots (a) and (b) show selected data sets from the Fig. reffig:cumuls2 (a,b), presented on the log-log scale to enhance the short-time dynamical range. In both plots, the power-law slopes of the short-time data follow the equation: α*t*^5^, with the coefficient prefactor *α* depending both on cell type and on temperature. The dashed line illustrate the slopes of *t*^6^ and *t*^4^ to illustrate the strength of fit.

However, the strong temperature dependence is evident in the short-time onset of mechanosensing: the difference was evident in Figs 2 and 3, but is very clearly enhanced in Fig. 5. What changes between the data sets is the prefactor *α* of the universal power law α *t*^5^, which has a systematic temperature dependence (the fitted values of α(*T*) are listed in the Supplementary). Now expecting the thermally activated behavior, by analogy with the earlier analysis, we plot these prefactors α(*T*) on the Arrhenius plot in Fig. 6. The fitting to *α* = const *e^−∆H/kB T^* indeed gives a very reasonable trend, with the activation energies ∆*H* = 70 kcal/mol for 3T3, and 129 kcal/mol for EA. Note that, in contrast to Fig. 4, here we have a negative exponent, i.e. the parameter α(*T*) represents a reaction *rate* rather than a relaxation *time*. In the classical Arrhenius-Kramers thermal activation, the process time is shorter as the temperature increases, while the Fig. 6 shows the scaling factor α(*T*) is decreasing as the temperature decreases instead (which is reflected in the overall observation of longer lag time in the cumulative curves). The magnitudes, and the difference in the energy barriers between the Biophysical Journal 00(00) 1–10 to transform these mechanical forces into a chemical signal, which is then starting one or several signaling pathways in the cell that lead to a morphological response (35–37): usually the enhancement of cytoskeleton by increasing the number and branching of F-actin, and activating more myosin motors. It is relatively well accepted that FAK is the central two cell types make sense because, due to their biological function, the mechanosensing process in fibroblasts should start faster. However, we have so many different quantitative facts and trends that it is necessary to look much more carefully at what we understand about mechanosensing.

**Figure 6:**
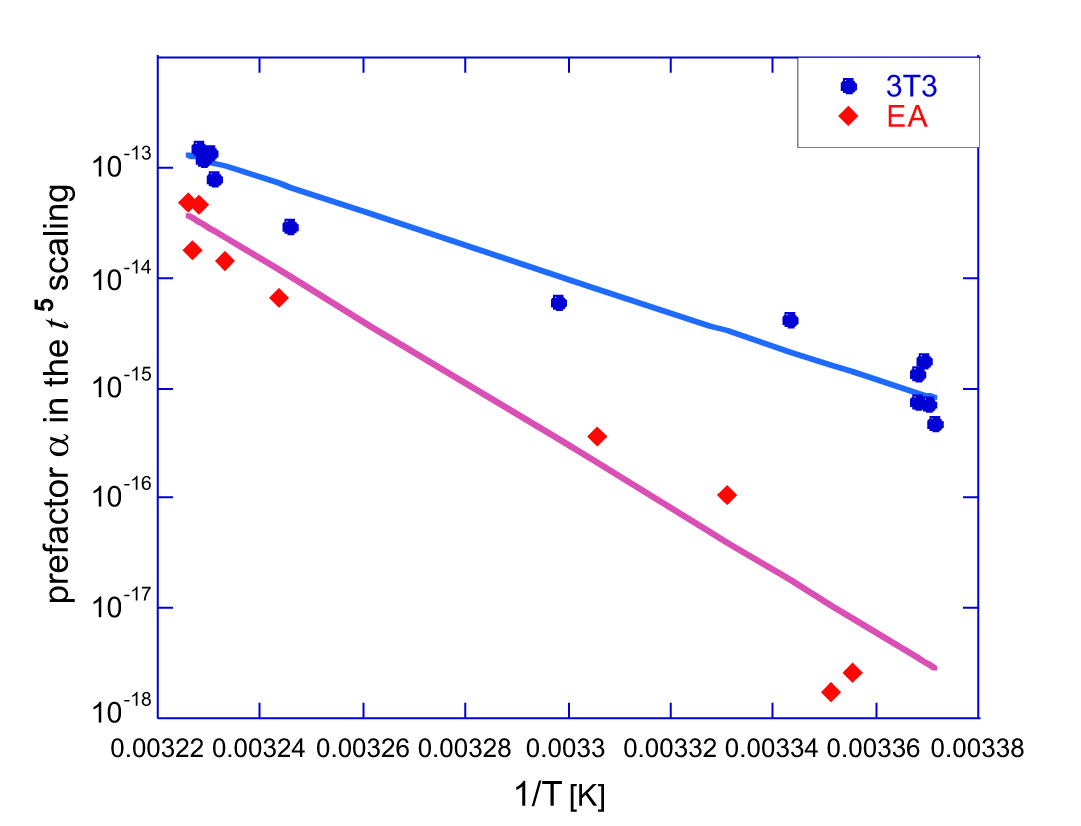
Analysis of the short-time dynamics of cell spreading. The Arrhenius plot of the prefactor α(*T*), with the fit lines giving the effective activation enthalpy ∆*H* ≈ 70 kcal/mol for 3T3, and 129 kcal/mol for EA. See text, explaining how this value represents the sum of free energy barriers of key proteins assembling into the adhesion-mechanosensing complex.

## 4 Discussion

To have such a mechanosensor operational, the cell needs to assemble the protein complex connecting and transmitting the pulling force from the F-actin terminals in the cytosol to the extracellular matrix (ECM). The mechanosensor has Biophysical Journal 00(00) 1–10 player in this signaling network (14, 35, 38, 39). A recent model of FAK as a mechanosensor (15) shows how the rate of its activation is sensitive to the stiffness of substrate, and the cytoskeletal pulling force, see some details in the Supplementary Information. Importantly, when the force is low (as we would expect at early times, before the mechanosensing pathways are activated and the cytoskeletal forces increase), this rate is controlled only by the bonding energy between its FERM and kinase domains, not the stiffness. A recent molecular-dynamics simulation (34) has explicitly calculated this bonding energy as ∆*G* ≈ 17 kcal/mol. If we associate this barrier with the longest relaxation time examined in Fig. 4, the agreement of the ∆*G* values is remarkably close. It makes sense that the mechanosensor itself should be the rate-limiting step of any signaling pathway!

What is the physical process underlying the apparently universal power law α*t*^5^ at the early stages of mechanosensing activation? Although previous work on cell spreading has also talked about power law behavior (19, 21, 23), it is important not to confuse the role of viscous dissipation in a spreading cell with the overall stochastic population dynamics of the onset of spreading. We believe the origin of this universal time dependence is the dynamic assembly of the integrin-FAK protein complex that is required to transmit the cytoskeletal pulling force across the cell membrane to the ECM bonding site (14), to initiate the mechanosensing response, see Figure 7. There is a large literature on all aspects of this complex composition and assembly (41, 42), with a consensus emerging in the recent years that the dimer of αβ-integrins needs to be activated to expose the ECM-binding ligand. This activation occurs on binding the N-terminal of talin (33, 43), while the C-terminal of talin is associated with paxillin, which in turn may associate with the focal adhesion targeting (FAT) domain (the C-terminal) of FAK. Both talin and paxillin also bind to cytoskeletal F-actin; talin-actin links are known to be strengthened by vinculin (44, 45), but we see in the sequence of steps illustrated in Fig. 7 that there is no separate distinct step of complex assembly required to account for that.

**Figure 7:**
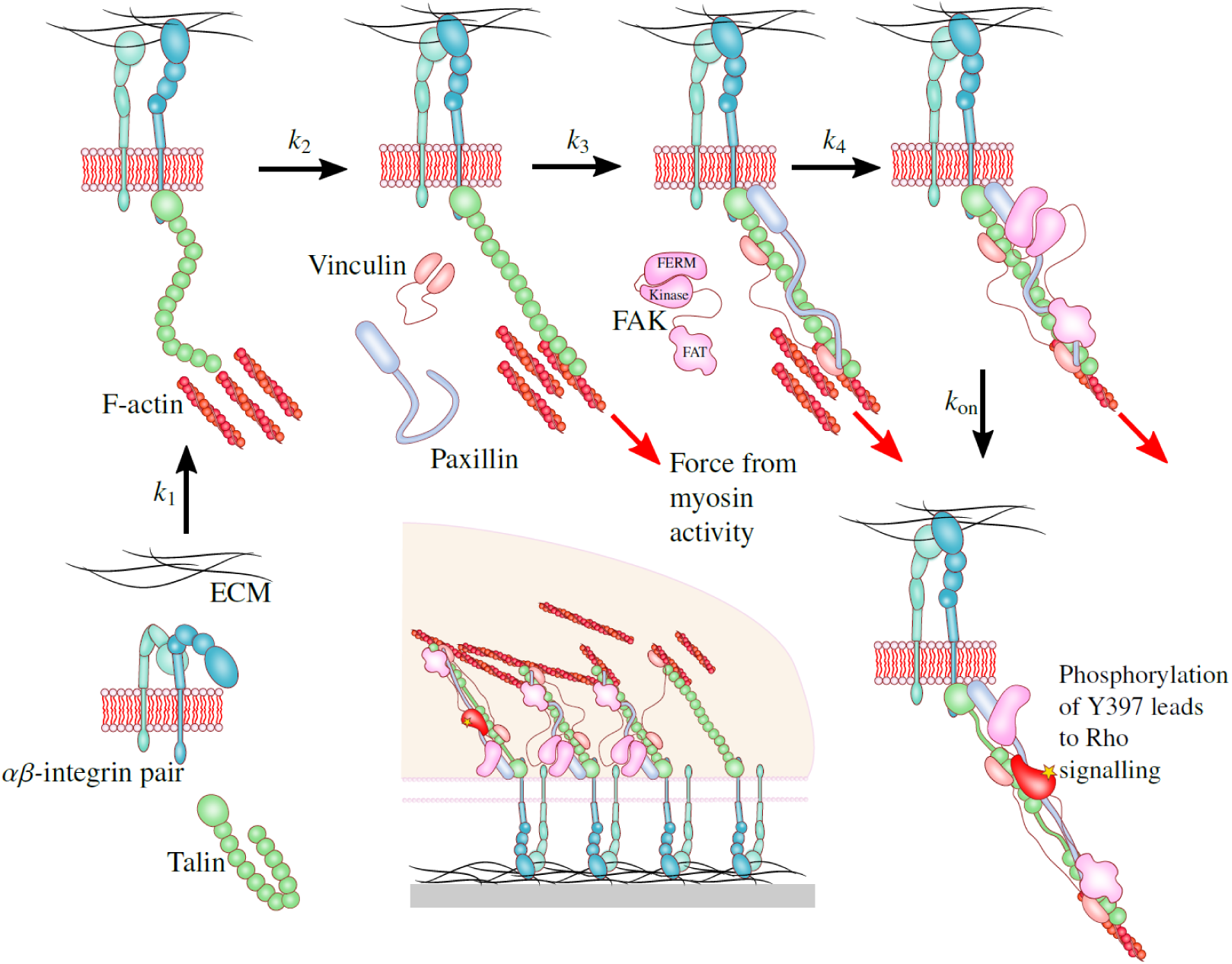
Assembly of a mechanosensor complex. Our analysis suggests that there are five distinct slow stages illustrated in the sequence, with their respective rates *k*_1_-*k*_4_ and the rate of FAK activation *k*_on_ (controlled by the free energy barrier *∆G* ≈ 19 kcal/mol, cf. Fig. 4). The product of the five rate constants *α* = *k*_1_*k*_2_*k*_3_*k*_4_*k*_on_ is what we measure in the Arrhenius plot Fig. 6. In the center is a sketch of forming focal adhesion cluster, where the individual mechanosensor complexes in various stages of development/turnover are bound by vinculin and actin crosslinking (40).

It is worth noting that while in well-developed adhesions there is considerable actin-myosin contraction, it may well be that in early stage spreading, instead of grabbing onto an actin filament and being subject to a myosin pulling force, the nascent complex instead captures actin filaments undergoing retrograde flow. The captured actin flow could start to force the leading edge out, and help activate FAK. It seems likely that FAK activation is needed for the cell to properly recruit actin to push out the leading edge; FAK-null cells are much slower in the stage of early spreading (46).

If we then summarize the suggested four steps of the essential complex assembly, they are the following: [1] Nterminal of talin binds and activates integrins; [2] integrins bind to ECM matrix, immobilizing the nascent complex, and C-terminal of talin binds to F-actin, thus forming the continuous force chain from cytoskeleton to ECM; [3] paxillin and/or vinculin bind to talin-actin assembly to complete the mechanical scaffold; [4] FAK binds to C-terminal of talin (and F-actin) by its FAT domain, and to the N-terminal of talin (and integrins) by its FERM domain. The sensor is now primed for action. The last step in the sequence is the opening of FAK domains, exposing the phosphorylation residues and the site for Src binding: the associate rate of FAK activation was calculated in (15).

The Supplementary Information gives more details of the classical kinetic theory (47) that predicts the early-time increase in the number of self-assembled nuclei of size *n* to grow as a power-law law α*t^n−^*^1^, with the prefactor α proportional to the product of binding rates of each contributing protein. The reason for this power-law is that each individual assembly state has a linear time dependence (at early times), and the total rate is the product of them. This kinetic theory of protein aggregation is an example of a more general network theory (48) that shows how the kinetics of complex pathways is determined by their topology. The observed short-time behavior leads us to predict that there are four slow processes in the integrin-FAK mechanosensor assembly (thus giving *t*^4^ growth), followed by the FAK activation process, completing the observed *t*^5^ law. By ‘slow’, we mean any processes with rates of *k <* 1min−^1^, because this was close to the resolution limit of our experiments; if a more advanced measurement of the kinetic curves such as in Fig. 5 could obtain data for the much shorter times after cell planting (say, within a second, or even sooner), this would resolve more fast assembly steps – and result in a steeper powerlaw in that very-short time region of the log-log plot. For instance, the dimerization of integrins, or the separate stages of talin binding and ECM binding, or other chaperon proteins attaching or assisting the complex construction could be resolved. But at our level of accuracy, the observed α *t*^5^ law leads to the 5-step sequence in Fig. 7.

Since the last step in our mechanosensing process has to be the FAK activation (the chemical signal generation), there are exactly four remaining kinetic steps left. That is, from the kinetic analysis we conclude that the sensor is an assembly of *n* = 5 essential proteins (via four steps of binding, with the aggregate rate *k*_1_*k*_2_*k*_3_*k*_4_). We may not be certain of the order of assembly (e.g. talin-actin first, or paxillin-talin), and we may not be certain about which proteins are essential (although integrins, talin, FAK and F-actin are certain) – but we have to be sure that there are exactly four steps of the complex assembly. The product of binding rates in the prefactor α gives the overall Arrhenius exponential we saw in Fig. 6 with the total activation energy being the sum of all processes. It appears that the sum of binding energies of integrin-ECM, talin-integrin, talin-FAK, and FAK-actin is different in cells with different biological function: 3T3 fibroblasts have this sum (70 – ∆*G*_FAK_) ≈ 51 kcal/mol, while EA endothelial cells have (129 – ∆*G*_FAK_) ≈ 110 kcal/mol.

In this work, we have studied the population dynamics of spreading cells, and from this macroscopic observation we were able to infer details of the microscopic processes governing the overall cellular response to an external substrate. By linking the results to nucleation theory, we have found a novel way of looking at the onset of cell spreading as a problem of complex assembly. The next step will be to understand the development of mature structures once the cell spreading is well underway.

## Author contribution

ALR carried out all experiments. SB, ALR and EMT carried out different elements of data analysis. SB and EMT wrote the paper.

## Acknowledgements

The authors acknowledge many helpful discussions with K. Franze, K. Chalut, A. B. Kolomeisky, and H. Welch. Critical comments of U. S. Schwarz, and experimental support in the cell culture lab by E. Nugent and F. Morgan are appreciated. This work has been funded by EPSRC (grants EP/M508007/1 and EP/J017639), and the Ernest Oppenheimer Trust in Cambridge.

